# BAZ2A association with H3K14ac is required for the transition of prostate cancer cells into a cancer stem-like state

**DOI:** 10.1101/2020.07.03.185843

**Authors:** Rodrigo Peña-Hernández, Rossana Aprigliano, Sandra Frommel, Karolina Pietrzak, Seraina Steiger, Marcin Roganowicz, Juliana Bizzarro, Raffaella Santoro

## Abstract

Prostate cancer (PCa) is one of the most prevalent cancers in men. Cancer stem cells are thought to be associated with PCa relapse. Here we show that BAZ2A is required for the transition of PCa cells into a cancer stem-like state. BAZ2A genomic occupancy in PCa cells coincides with H3K14ac enriched chromatin regions. This association is mediated by BAZ2A-bromodomain (BAZ2A-BRD) that specifically binds H3K14ac. BAZ2A associates with inactive enhancers marked by H3K14ac and repressing transcription of genes frequently silenced in aggressive and poorly differentiated PCa. BAZ2A-mediated repression is also linked to EP300 that acetylates H3K14ac. BAZ2A-BRD mutations or treatment with inhibitors abrogating BAZ2A-BRD/H3K14ac interaction impair the transition of PCa cells into a stem-like state. Furthermore, pharmacological inactivation of BAZ2A-BRD impairs *Pten*-loss oncogenic transformation of prostate organoids. Our findings indicate a role of BAZ2A-BRD in PCa stem cell features and suggest potential epigenetic-reader therapeutic strategies to target BAZ2A in aggressive PCa.

## Introduction

Tumors are composed of heterogeneous populations of cells, which differ in their phenotypic and genetic features (Saygin et al., 2019). It has been suggested that cellular heterogeneity within a tumor is organized in hierarchical manner with a subpopulation of cancer stem cells (CSC) that give rise to heterogeneous cancer cell lineages and undergo self-renewal to maintain their reservoir (Pattabiraman and Weinberg, 2014). Furthermore, non-stem, differentiated cancer cells can transit into a dedifferentiated, CSC-like phenotype (Plaks et al., 2015; Rich, 2016). CSCs were shown to have an enhanced capacity for therapeutic resistance, immune evasion, invasion, and metastasis (Prager et al., 2019). Thus, efficient targeting of CSCs in cancer treatment is critical for developing effective therapeutics.

Prostate cancer (PCa) is the second most common epithelial cancer and the fifth leading cause of cancer-related death in men worldwide (Bray et al., 2018). PCa displays a high heterogeneity that leads to distinct histopathological and molecular features (Li and Shen, 2018). This heterogeneity posits one of the most confounding and complex factors underlying its diagnosis, prognosis, and treatment (Yadav et al., 2018). Current therapeutic approaches for PCa remain insufficient for some patients with progressive disease. Targeting of the androgen receptor (AR) axis through androgen ablation therapy and/or AR antagonists (castration) in patients with PCa relapse was shown to be efficient only for a short period of time (Denmeade and Isaacs, 2002). These treatments are not curative; a population of cells resistant to androgen-deprivation therapy emerges and PCa becomes unresponsive and progresses to a castrate-resistant prostate cancer (CRPC) and metastasis, with limited treatment options (Linder et al., 2018). Due to the enticing possibility that PCa aggressiveness and relapse arises from PCa stem cells (Colombel et al., 2012; Mayer et al., 2015; Seiler et al., 2013), the development of stem cell-specific anticancer drugs constitutes an attempt to innovate in the treatment of PCa (Leao et al., 2017).

Targeting of bromodomain (BRD)-containing proteins is a promising therapeutic approach, currently undergoing clinical evaluation in CRPC patients (Welti et al., 2018). BRD-containing proteins regulate gene expression primarily through recognition of histone acetyl residues, leading to the recruitment of protein complexes that modulate gene expression (Fujisawa and Filippakopoulos, 2017). BRD containing proteins are frequently deregulated in cancer and the development of BRD inhibitors as anticancer agents is now an intense area of research (Fujisawa and Filippakopoulos, 2017). JQ1, an inhibitor of the BET family of BRDs has an effect in AR-signaling-competent human CRPC cell lines, whereas in AR-independent cell lines, such as PC3 cells, these inhibitors do not display any effect (Asangani et al., 2014).

BAZ2A (also known as TIP5) is a BRD-containing protein and subunit of the nucleolar-remodeling complex NoRC (Santoro et al., 2002; Zhou and Grummt, 2005). Recent work implicated BAZ2A in aggressive PCa. BAZ2A is highly expressed in metastatic tumors compared to primary and localized tumors, required for cell proliferation, viability, and invasion, and represses genes frequently silenced in aggressive PCa (Gu et al., 2015). Furthermore, high BAZ2A levels in tumors associate with poor prognosis and disease recurrence. Finally, it was shown that BAZ2A is required for the initiation of PCa driven by ***PTEN*-**loss, one of the most commonly lost tumor suppressor gene in PCa (Pietrzak et al., 2020).

BAZ2A contains a BRD that binds to acetylated histones (Tallant et al., 2015; Zhou and Grummt, 2005). In this work, we show that BAZ2A genomic occupancy in PCa cells coincides with H3K14ac enriched chromatin regions. This association is mediated by BAZ2A-BRD, an epigenetic reader of H3K14ac. In PCa cells, BAZ2A-BRD is required for the interaction with a class of inactive enhancers that are marked by H3K14ac and represses the expression of genes implicated in developmental and differentiation processes that are linked to aggressive and dedifferentiated PCa. BAZ2A-mediated repression of these genes is also linked to the histone acetyltransferase EP300 that acetylates H3K14ac and displays the highest positive correlation with BAZ2A expression in both metastatic and primary tumors compared to the other H3K14 acetyltransferases. Mutations of BAZ2A-BRD or treatment with chemical probes that abrogate BAZ2A-BRD association with H3K14ac impair the transition of PCa cells into a stem-like state. Furthermore, pharmacological inactivation of BAZ2A-BRD impairs the oncogenic transformation mediated by *Pten*-loss in PCa organoids. Our findings indicate that BAZ2A is a key player in PCa stem cell features and suggest potential epigenetic-reader therapeutic strategies to target BAZ2A-BRD in aggressive, poorly differentiated PCa.

## Results

### BAZ2A associates with chromatin regions marked by H3K14ac

To determine whether BAZ2A-BRD domain is implicated in BAZ2A-mediated regulation of PCa, we investigated its genomic occupancy and the correlation with histone marks in metastatic PCa PC3 cells. We performed ChIPseq analysis of PC3 cells transfected with plasmids expressing HA-tagged BAZ2A and found 96351 regions that were bound by BAZ2A (**Fig. 1a**). We validated these results by ChIP-qPCR through the measurement of the binding of endogenous BAZ2A with two selected genes (*AOX1*, that as described below is a gene directly regulated by BAZ2A, and *EVX2*) using a PC3 cell line that expresses endogenous BAZ2A with a FLAG-HA (F/H) tag at the N-terminus (**Fig. 1b,c and Supplementary Fig. 1a,b**). The correlation between ChIPseq signal enrichment for BAZ2A binding and histone marks in PC3 cells indicated that BAZ2A has the strongest association with H3K14ac whereas its interaction with H3K27ac regions was much weaker (**Fig. 1a,b,d**). To determine whether BAZ2A-BRD specifically recognizes H3K14ac, we used histone peptide arrays containing 384 unique histone modification combinations that allow the analysis of not only individual modified residues but also the effects of neighboring modifications (**Figure 1e, Supplementary Fig. 1c**). We incubated purified recombinant human BAZ2A-BRD fused to GST-tag with the histone array and monitored the binding to histone peptides using anti-GST antibodies. Consistent with previous results (Tallant et al., 2015; Zhou and Grummt, 2005), BAZ2A-BRD did not interact with unmodified histone peptides and showed a preferential binding to a specific set of acetylated histone peptides. In line with the ChIPseq analyses, the acetylation per se was not sufficient for the binding of BAZ2A-BRD as evident by the lack of interaction with acetylated histone peptides such as H3K27ac. Interestingly, H3K14ac peptide was the mono-acetylated histone peptide displaying the highest interaction with BAZ2A-BRD, a results that is consistent with the ChIPseq analysis. Phosphorylation of H3S19 or acetylation of H3K9 improved the interaction of BAZ2A with H3K14ac. Poly-acetylated histone H4 containing K5ac, K8ac, K12ac and K16ac was also a good target for BAZ2A-BRD whereas the interaction with these single H4 acetylated residues was much weaker or even absent as in the case of H4K5ac. To further support the specificity of BAZ2A-BRD as reader of H3K14ac, we mutated BAZ2A-Y1830 and N1873, which recent structural analyses indicated as critical residues for the interaction of BAZ2A-BRD with acetylated histones (Tallant et al., 2015). Accordingly, the interaction of both BAZ2A-BRD_Y1830F_ and BAZ2A-BRD_N1873L_ with modified histone peptides was strongly reduced, particularly for the mono-acetylated H3K14ac peptide (**Fig. 1e**).

**Figure 1.**
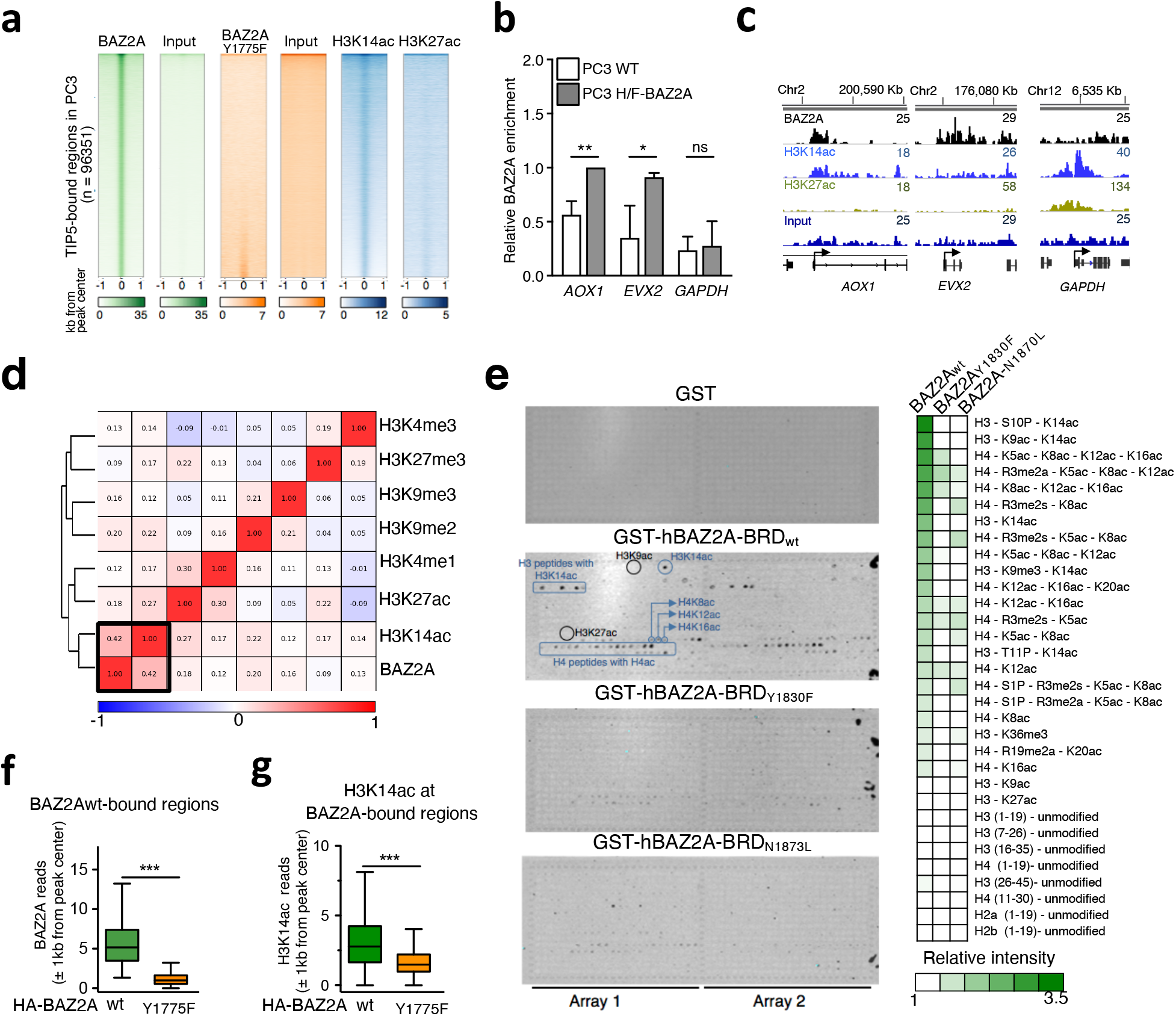
BAZ2A-BRD is an epigenetic reader of H3K14ac that mediates the BAZ2A association with H3K14ac-chromatin domains in PC3 cells. **a**. Heatmap profiles of BAZ2A bound regions in PC3 cells and the corresponding signals of input, BAZ2A-BRD mutant (Y1775F), H3K14ac and H3K27ac. Data were ranked to BAZ2A-binding in PC3 cells. **b**. Validation of ChIPseq results by anti-HA ChIP-qPCR of PC3 wild-type (WT) and a PC3 cell line that expresses endogenous BAZ2A with a FLAG-HA tag (PC3-H/F-BAZ2A). *GAPDH* promoter represents a region not bound by BAZ2A. Average values of three independent experiments. Data were normalized to input and to AOX1 levels. Statistical significance (*P*-values) was calculated using two-tailed t-test (*<0.05, **< 0.001, ns: not significant) **c**. Wiggle tracks displaying BAZ2A-bound regions, H3K14ac, H3K27ac, and input at *AOX1, EVX2*, and *GAPDH*. **d**. Pearson correlation heat map of BAZ2A-bound regions and histone marks. Data for BAZ2A, H3K4me3, H3K27me3, H3K27ac, H3K14ac are from this work. H3K9me3, H3K9me2, H3K4me1 were obtained from ENCODE. **e**. Images of histone peptide arrays probed with 10nM of recombinant BAZ2A-BRD wild-type (GST-BAZ2A-BRD_wt_) and mutants (GST-BAZ2A-BRD_Y1830F_ or GST-BAZ2A-BRD_N1873L_). Visualization of binding was performed by incubation with anti-GST antibodies and imaged on Odyssey Infrared Imaging System. Blue circles mark peptides recognized by BAZ2A-BRD (i.e. H3K14ac), whereas black circles show some of the acetylated peptides not recognized by BAZ2A-BRD (H3K27ac and H3K9ac). Right panel shows heatmap of the relative binding intensity of BAZ2A-BRD against modified histone peptides. Binding intensity was calculated as average of fold change from peptides signal over background controls of 2 different arrays. **f**. BAZ2A read coverage quantification at ±1Kb from peak summit of BAZ2A-bound regions identified in PC3 cells and the corresponding reads for BAZ2A-BRD mutant (Y1775F) in PC3 cells. Statistical significance (*P*-values) was calculated using two-tailed t-test (***< 0.001). **g**. Regions bound by BAZ2A in PC3 cells are enriched in H3K14ac. Read coverage quantification for H3K14ac levels at regions bound by BAZ2A in PC3 cells (BAZ2Awt and BAZ2A-BRD mutant (Y1775F)) at ±1Kb from BAZ2A peak summits. Statistical significance (*P*-values) was calculated using two-tailed t-test (***< 0.001).

To determine whether BAZ2A-BRD mediates the association of BAZ2A with chromatin, we performed ChIPseq analysis of PC3 cells expressing murine BAZ2A-BRD mutant HA-BAZ2A_Y1775F_, an orthologous to the human BAZ2A-BRD_Y1830F_ unable to bind to acetylated histones (Zhou and Grummt, 2005). Equal expression of HA-BAZ2Awt and HA-BAZ2A_Y1775F_, was assessed by western blot (**Supplementary Fig. 1d**). We found that only 1.3% (1534 peaks) of BAZ2A-bound sites were also occupied by BAZ2A_Y1775F_, indicating that a functional BRD domain is required for BAZ2A binding to chromatin in PC3 cells **Fig. 1a,f**). Furthermore, the few DNA regions that retained binding with BAZ2A-BRD mutant contained low H3K14ac levels (**Fig. 1g**). Together, these results suggest that BAZ2A is an epigenetic reader of H3K14ac. Moreover, they show that in PC3 cells BAZ2A preferentially binds to regions marked by H3K14ac and that this interaction is mediated by BAZ2A-BRD.

### BAZ2A binds to inactive enhancers enriched in H3K14ac and mediates the repression of genes linked to developmental and differentiation processes

Previous studies showed that acetylation of H3K14 is mediated by histone acetyltransferases (HATs) EP300, KAT2A (GCN5), or KAT6A (Myst3) (Jin et al., 2011; Lee and Workman, 2007). Interestingly, we found that all these HATs are highly expressed in metastatic prostate cancer (**Supplementary Fig. 2a**). In PC3 cells, however, only EP300 and KAT6A are expressed whereas *KAT2A* sequences are deleted (Seim et al., 2017) (**Supplementary Fig. 2b**). The expression profile of these HATs in a large cohort of primary PCa and metastatic CRPC (Cancer Genome Atlas Research, 2015; Robinson et al., 2015) revealed that the levels of EP300 had the highest positive correlation with BAZ2A expression in both metastatic and primary tumors compared to KAT2A and KAT6A, suggesting that EP300 might be functionally related with BAZ2A binding to H3K14ac chromatin and the regulation of gene expression in PCa (**Fig. 2a, Supplementary Fig. 2c**). Treatment of PC3 cells with A-485, a selective catalytic EP300 inhibitor (Lasko et al., 2017), induced a decrease of H3K14ac levels, confirming the role of EP300 in acetylating H3K14 (**Supplementary Fig. 2d**). To test the functional link between BAZ2A and EP300, we initially performed genomic annotation analyses of the BAZ2A-bound sites that contain H3K14ac in PC3 cells. We found that BAZ2A is mainly located in intronic or intergenic regions, whereas only a small fraction (11%) was located at gene promoters (**Fig. 2b**). To determine which of the genes with BAZ2A-bound promoter transcriptionally depend on BAZ2A, we performed RNAseq of PC3 cells depleted of BAZ2A by siRNA. Consistent with previous results (Gu et al., 2015), BAZ2A-KD affects gene expression in PC3 cells (log_2_ fold change > 1; *P*<0.05; 535 upregulated genes, 819 downregulated genes) (**Supplementary Fig. 2e**). However, we found that only a minority of genes with BAZ2A-bound promoter was transcriptionally affected upon BAZ2A depletion (30 upregulated and 25 downregulated out of 527 genes). Although the interaction of BAZ2A with the promoter does not appear the most prominent mechanism for BAZ2A-mediated regulation in PC3 cells, we found that depletion of EP300 by siRNA reactivated the expression of *AOX1, DDB1, MANB2*, and *DMPK* (**Supplementary Fig. 2f,g**). Interestingly, these are the only upregulated genes upon BAZ2A-KD with BAZ2A-bound promoter that are significantly lower expressed in metastatic PCa compared to normal prostate tissue (**Supplementary Fig. 2f,h**). Furthermore, low expression of *AOX1* has recently been correlated with shorter time to biochemical recurrence of PCa (Li et al., 2018). These results indicate that EP300 is required for BAZ2A-mediated repression of a set of genes with BAZ2A-bound promoter that are implicated in metastatic PCa.

Given that a large fraction of BAZ2A-bound regions are intronic and intergenic, we asked whether BAZ2A could regulate gene expression through its association with enhancers. First, we classified annotated enhancers (Plaschkes et al., 2017) at intergenic regions according to histone marks and chromatin accessibility and found that a large fraction of H3K14ac peaks in PC3 cells are located at active enhancers marked by H3K27ac, H3K4me1, and DNase I hypersensitive sites (from here on named as enhancer class 1, C1, **Fig. 2c**). However, this class of active enhancers was not bound by BAZ2A. Interestingly, we found that BAZ2A only associates with a class of enhancers (named as C2) that relative to C1 enhancers had higher H3K14ac and H3K27me3 levels, lower content of H3K27ac, and lacked H3K4me1 and DNase I hypersensitive sites (**Fig. 2c,d**). These results suggest that in PC3 cells BAZ2A binds to a class of inactive enhancers that is enriched in H3K14ac. To determine whether the binding of BAZ2A to C2-enhancers affects gene expression, we analyzed the expression levels of genes in the nearest linear proximity to BAZ2A-bound C2-enhancers upon BAZ2A-KD in PC3 cells (**Fig. 2e,f**). We found that 56% of these genes (417 out of 747) were significantly affected in their expression upon BAZ2A depletion. Furthermore, the majority of these BAZ2A-regulated genes (254 genes, 61%) showed upregulation upon BAZ2A-KD, suggesting a major role of BAZ2A in repressing gene transcription through its interaction with C2-enhancers. Interestingly, the top 10 gene ontology (GO) terms of these BAZ2A-regulated genes were strongly linked to developmental and differentiation processes (**Fig. 2g**). Histological grading from well-differentiated to poorly differentiated PCa measured by Gleason score marks the progression from low to high-grade cancer (Ahmed et al., 2012). We set to identify BAZ2A-bound C2-enhancer regulated genes that were upregulated upon BAZ2A-KD and lower expressed in Gleason 9 and 10 tumours (advanced) compared to Gleason 6 tumours (indolent). Using these criteria, we found 25 genes (**Fig. 2h and Supplementary Fig. 2i**). Since GO terms associated with these genes were linked to developmental and cellular growth pathways, we selected five genes (*KRT8, EVC, ITPKB, SH3BP4*, and *SUN2*) that were implicated in these processes and analysed their expression upon EP300 depletion in PC3 cells (**Fig. 2i,j, Supplementary Fig. 2g**). We found that EP300-KD increased the expression of all the selected genes, a result that further supports the functional link between BAZ2A and EP300 in the regulation of gene expression in PCa cells. Collectively, these results suggest that BAZ2A binds to inactive C2-enhancers containing H3K14ac and mediates the repression of genes linked to developmental and differentiation processes that are frequently silenced in aggressive and poorly differentiated tumors.

**Figure 2.**
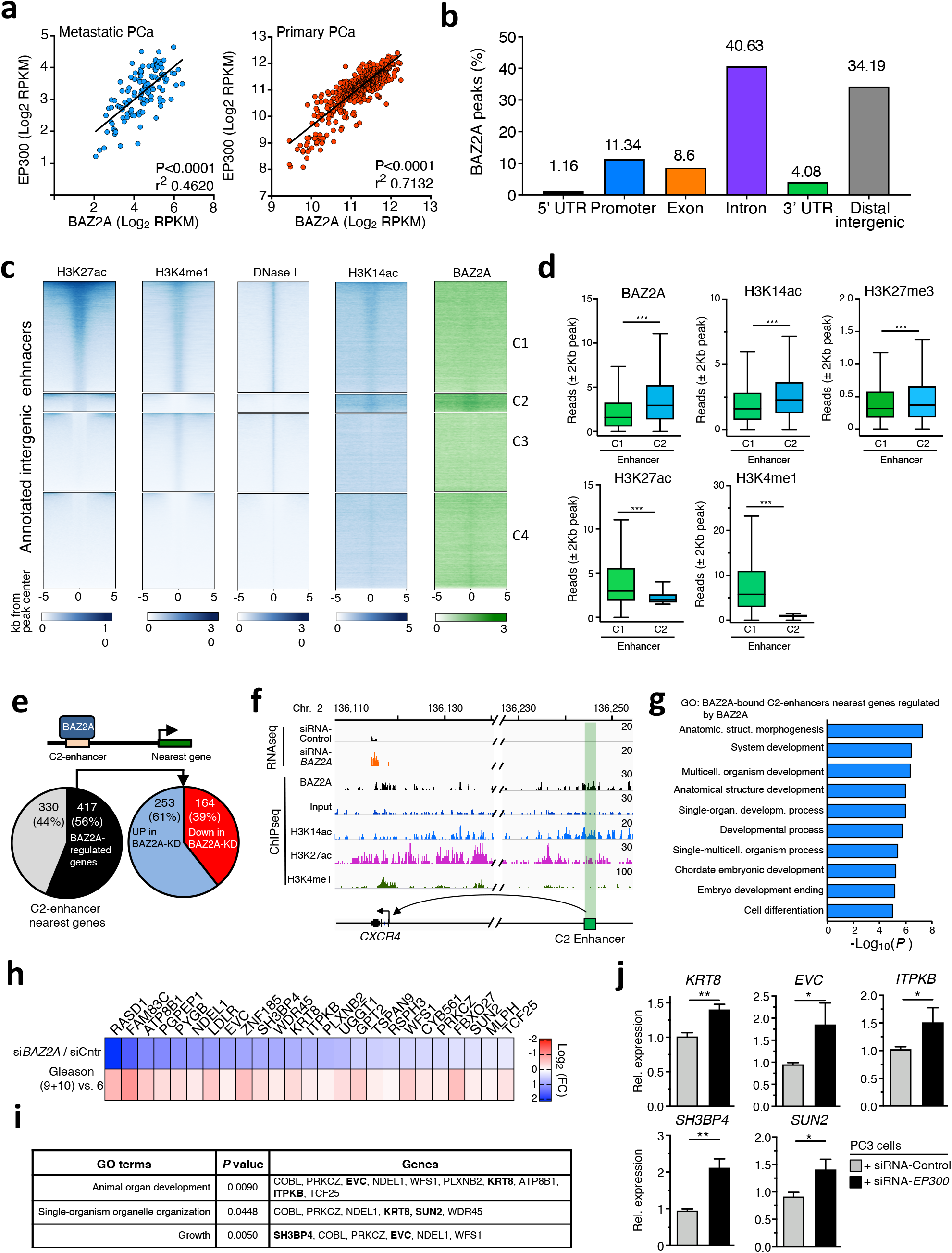
BAZ2A binds a class of inactive enhancer containing H3K14ac. **a**. Scatter plot showing the expression of *EP300* and *BAZ2A* in two large cohorts of metastatic and primary PCa. Data were from (Cancer Genome Atlas Research, 2015; Robinson et al., 2015). **b**. Genomic annotations of BAZ2A bound regions in PC3 cells. **c**. BAZ2A binds to a class of inactive enhancers marked by H3K14ac. Heat maps of BAZ2A, H3K14ac, H3K27ac, H3K4me1, and DNAseq signal at annotated intergenic enhancers are shown. Enhancer regions were clustered in 4 groups based on the presence (+) or absence (-) of H3K27ac and H3K4me1. C1 (H3K27ac+ / H3K4me1+), C2 (H3K27ac+ / H3K4me1-), C3 (H3K27ac- / H3K4me1+), and C4 (H3K27ac- / H3K4me1-). **d**. Inactive enhancers (C2) are enriched in H3K14ac and BAZ2A levels. Read coverage at enhancer classes C1 and C2 for BAZ2A, H3K14ac, H3K27ac, H3K4me1 and H3K27me3 at ±1Kb from annotated enhancer regions. Statistical significance (P-values) was calculated using two-tailed t-test (***< 0.001). **e**. Pie charts showing the number of genes in the nearest linear genome proximity to BAZ2A-bound C2 enhancers and their expression changes upon BAZ2A-KD in PC3 cells. **f**. Wiggle tracks showing RNA levels in PC3 cells treated with siRNA-BAZ2A and occupancy of BAZ2A, H3K14ac, H3K27ac, and H3K4me1 at C2 enhancer and its nearest gene *CXCR4*. **g**. Top ten biological process gene ontology (GO) terms as determined using DAVID 6.8 for genes in the nearest linear proximity to BAZ2A-bound C2-enhancers that are differentially expressed upon BAZ2A-KD in PC3 cells. **h**. Heatmap showing expression changes of BAZ2A-bound C2-enhancer genes that were both significantly upregulated upon BAZ2A-KD and lower expressed in Gleason 9 and 10 tumours (advanced) compared to Gleason 6 tumours (indolent). Values from tumours were from (Cancer Genome Atlas Research, 2015). **i**. GO terms associated with the genes listed in **H**. Genes labeled in bold were analyzed in **J**. **j**. qRT-PCR showing increased expression levels of BAZ2A-bound C2-enhancer genes in PC3 cells treated with siRNA-EP300. Data were from three independent experiments. Downregulation of *EP300* expression is shown in **Fig. S2G**. Values were normalized to *GAPDH* mRNA. Statistical significance (P-values) was calculated using two-tailed t-test (*<0.05, **<0.001).

### BAZ2A-BRD is required for PCa stem cells

Several studies have reported that non-stem, differentiated cancer cells can transit into a dedifferentiated, CSC-like phenotype (Plaks et al., 2015; Rich, 2016). The role of BAZ2A in modulating the expression of genes linked to developmental processes and frequently silenced in aggressive and poorly differentiated tumors prompted us to determine whether BAZ2A plays a role in the transition of cancer cells into a dedifferentiated, CSC-like state and whether this process is mediated by BAZ2A-BRD. To study this, we analyzed the capacity of PC3 cells to transit into CSC state by measuring the amounts of tumorspheres generated using serum-free medium and low attachment culture conditions (Li et al., 2008; Portillo-Lara and Alvarez, 2015; Rajasekhar et al., 2011a; Sheng et al., 2013). The PCa sphere assay recapitulates the ability of cancer cells to transit between stem and differentiated states in response to therapeutic insults or other stimuli within the microenvironment (Plaks et al., 2015; Rich, 2016) and has also been utilized in clinical trials testing anti-CSC therapeutics (Saygin et al., 2019). In particular, PC3 cells were shown to contain a small fraction (∼1%) of cells that can generate tumorspheres (Li et al., 2008; Sheng et al., 2013) and become significantly more clonogenic, chemoresistant, and invasive compared to the heterogeneous PC3 cell population (Civenni et al., 2013; Portillo-Lara and Alvarez, 2015). Tumorspheres isolated from PC3 cells (PC3-CSCs) could be serially passaged *in vitro* and when plated in adherent growth conditions they could differentiate into monolayer cultures morphologically indistinguishable from the original PC3 cells (**Supplementary Fig. 3a**). To further characterize the obtained PC3-CSCs, we performed RNAseq analysis and found that 1906 genes display transcriptional changes (log_2_ fold change > 1; *P* <0.05; 922 upregulated and 984 downregulated in PC3-CSCs vs. PC3 cells) (**Fig. 3a**). GO analysis indicated that genes downregulated in PC3-CSCs were associated with developmental processes (i.e. *HOXA1, HOXA4, SOX7*), reflecting a more dedifferentiated state compared to the parental PC3 cells (**Fig. 3b,c**). Furthermore, gene set enrichment analysis (GSEA) using predefined embryonic stem cell (ESC)-like gene signatures showed that genes upregulated in PC3-CSCs were enriched in adult stem cell signatures (Wong et al., 2008) and displayed a transcription profile similar to genes highly expressed in human embryonic stem cells compared to differentiated cells (Ben-Porath et al., 2008; Wong et al., 2008) (**Supplementary Fig. 3b**). Importantly, among the upregulated genes we found markers associated with CSCs such as *LGR5, CD24, Nestin, EpCAM* and *ALDH1A2* (Cao et al., 2017; Li et al., 2017; Raha et al., 2014) (**Fig. 3c, Supplementary Fig. 3c**). Taken together, these results indicated that tumorspheres derived from PC3 cells have a gene signature that resemble the molecular signature of a cancer stem cell-like phenotype.

**Figure 3.**
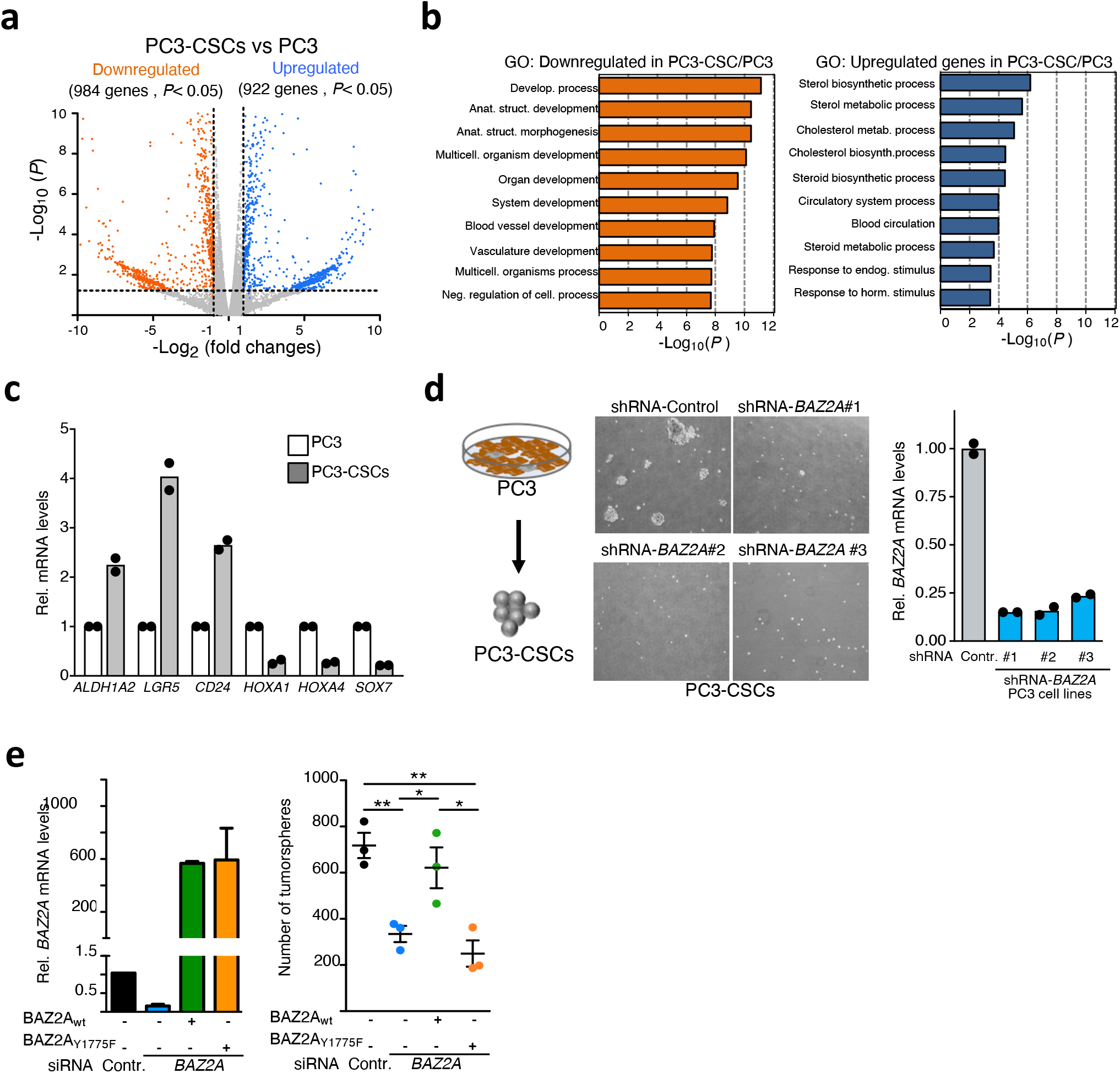
BAZ2A-BRD is required for the transition of PCa cells into a dedifferentiated, cancer stem-like state. **a**. RNA-seq showing differential gene expression between PC3-CSCs and the parental PC3 cells (P<0.05, Log_2_ fold change 1). **b**. Top ten biological process gene ontology (GO) terms as determined using DAVID 6.8 for genes downregulated and upregulated in PC3-CSCs relative to parental PC3 cells. **c**. qRT-PCR of genes differentially expressed in PC3-CSCs compared to PC3 cells showing the upregulation of CSC markers (*ALDH1A2, LGR5, CD24*) *and* downregulation of genes related to developmental processes (*HOXA1, HOXA4, SOX7*). Data are from two independent experiments. mRNA levels were normalized to *GAPDH*. **d**. BAZ2A is required for the dedifferentiation of PC3 into PC3-CSCs. Left panel: Representative images from two independent experiments showing the impairment of PC3 cells to form tumorspheres upon stable depletion of BAZ2A. Right panel: *BAZ2A* mRNA levels in three independent PC3 cell lines stably expressing shRNA-*BAZ2A* were measured by qRT-PCR and normalized to *GAPDH* mRNA from 2 independent experiments. **e**. Left panel: qRT-PCR showing *BAZ2A* mRNA levels in PC3 cells depleted of endogenous *BAZ2A* with siRNA specifically targeting human *BAZ2A* sequences and expressing mouse BAZ2Awt and BAZ2A-BRD mutant (BAZ2A_Y1775F_). Values were normalized to *MRPS7* mRNA. Right panel: Quantification of tumorspheres from 3 independent experiments. Statistical significance (*P*-values) was calculated using two-tailed t-test (* <0.05; **< 0.01).

To determine whether BAZ2A is implicated in the transition of cancer cells into a CSC-like phenotype, we established three PC3 cell lines stably expressing shRNA against *BAZ2A* sequence (**Fig. 3d**). Remarkably, all shRNA-*BAZ2A* PC3 cell lines were unable to generate PC3-CSCs. Similarly, treatment of PC3 cells with siRNA-*BAZ2A* reduced their ability to transit into CSCs (**Fig. 3e**). This phenotype could be rescued upon ectopic expression of mouse BAZ2A, which is not targeted by the siRNA-*BAZ2A*. Importantly, the expression of the BRD-mutant mBAZ2A_Y1775F_ abrogated the formation of tumorspheres, indicating that a functional BAZ2A-BRD is critical for the transition of PC3 cells into CSCs (**Fig. 3e**). Collectively, these results indicate that a functional BAZ2A-BRD is required for the transition of PC3 cells into CSCs and supported a role of BAZ2A in driving aggressive and poorly differentiated PCas.

### Pharmacological inhibition of BAZ2A-BRD impairs the transition of PCa cells into a stem-like state and the oncogenic transformation mediated by *Pten*-loss

The requirement of a functional BAZ2A-BRD for the transition of PC3 cells into CSCs, prompted us to test the possibility to inhibit BAZ2A-BRD for targeting CSCs as a therapeutic strategy. Recently, two chemical probes, GSK2801 and BAZ2-ICR, have been generated for specific targeting of BRDs of BAZ2A and the closely related BAZ2B (Chen et al., 2015; Drouin et al., 2015), which display 65% sequence similarity (Tallant et al., 2015). Both compounds are highly specific for BAZ2A-BRD and BAZ2B-BRD and show half-maximal inhibitory concentration (IC50) in the nanomolar range (Chen et al., 2015; Drouin et al., 2015). To determine whether BAZ2-ICR or GSK2801 destabilize BAZ2A-BRD interaction with histones, we incubated recombinant BAZ2A-BRD with both compounds and measured BAZ2A-BRD binding to modified histones with the histone peptide array (**Supplementary Fig. 4a,b**). Both BAZ2-ICR and GSK2801 abolished the interaction of BAZ2A-BRD with histones, although GSK2801 inhibition was weaker than BAZ2-ICR.

Next, we tested the possibility to impair BAZ2A function in PCa cells through pharmacological targeting of BAZ2A-BRD. Previous results showed that BAZ2A depletion in several metastatic PCa cells, including PC3 cells, impairs cell proliferation (**Fig. 4a**) (Gu et al., 2015). Indeed, all our attempts to isolate BAZ2A-KO PC3 cell lines through CRIPS/Cas9 failed, indicating the requirement of BAZ2A expression in metastatic PCa cells. Surprisingly, treatment of PC3 cells with BAZ2-ICR or GSK2801 at concentrations up to 50μM did not affect cell proliferation (**Fig. 4b, Supplementary Fig. 4c)**, suggesting that BAZ2A-BRD is not functionally relevant in the proliferation of the heterogeneous PC3 cell population. In contrast, treatment with BAZ2-ICR or GSK2801 drastically reduced the formation of tumorspheres (EC_50_ 0.83μM; **Fig. 4c,d**), indicating that pharmacological targeting of BAZ2A-BRD blocks the transition of PC3 cells into CSCs. The abrogation of tumorsphere formation upon treatment with BAZ2A-BRD inhibitors could also be observed in another metastatic PCa cell line, DU-145 (**Fig. 4e**). These results are consistent with the data showing the ability of PCa cells to transit into a CSC-like state requires a functional BAZ2A-BRD (**Fig. 3e**). Since BAZ2-ICR or GSK2801 can also target BAZ2B-BRD (Chen et al., 2015; Drouin et al., 2015), we tested the contribution of BAZ2B in the ability of PC3 cells to transit into CSCs by transfecting PC3 cells with siRNA directed against *BAZ2B*. We found no detectable defects compared to control cells (**Supplementary Fig. 4d**), indicating that pharmacological targeting of BAZ2A-BRD has a major function in the transition of PCa cells into a CSC-like state.

**Figure 4.**
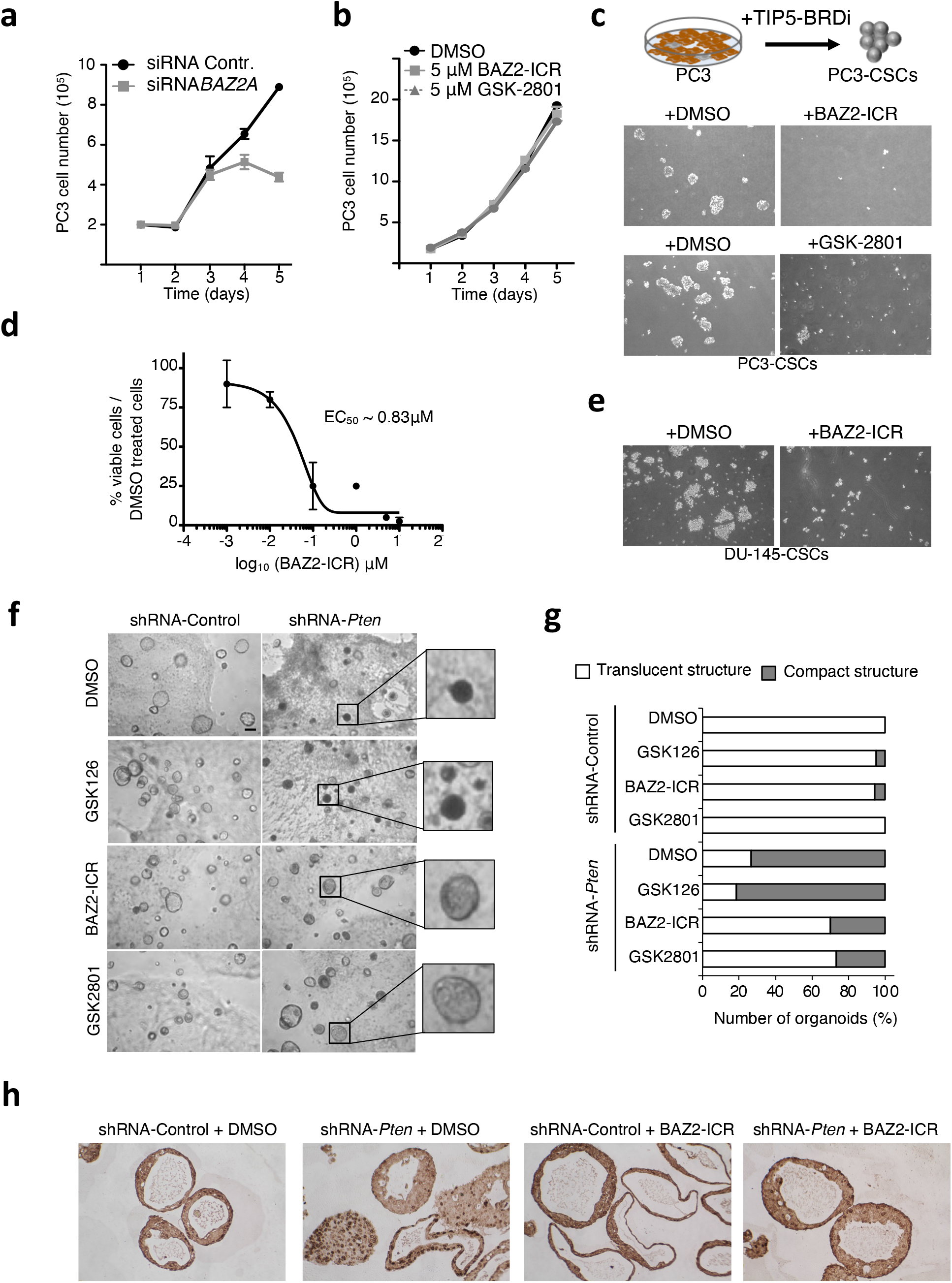
Pharmacological targeting of BAZ2A-BRD impairs the transition of PCa cells into a dedifferentiated, cancer stem-like state. **a**. Cell proliferation curves of PC3 cells treated with siRNA-*BAZ2A*. Error bars represent the standard deviation from three independent experiments. **b**. Cell proliferation curves of PC3 cells treated with 5μM BAZ2-ICR and GSK2801. Error bars represent the standard deviation from three independent experiments. **c**. Representative images of tumorspheres generated from PC3 cells treated with BAZ2-ICR (1μM) or GSK2801 (1μM). **d**. Half-maximal effective concentration (EC50) of BAZ2-ICR. Measurements were performed 5 days after treatment of PC3 cells and initiation of tumorspheres using the indicated BAZ2-ICR concentration. The amount of tumorspheres was assessed by measurement of cell viability. **e**. Representative images showing tumorspheres derived from the metastatic PCa cell line DU-145 treated with DMSO or BAZ2-ICR (1μM) inhibitor for 5 days. **f**. Representative bright field images of organoids derived from mouse prostate cells treated with DMSO, BAZ2-ICR (5μM), GSK2801 (5μM) or GSK126 (5μM) inhibitors followed by transduction with shRNA-Control or shRNA-*Pten*. Scale bar represents 200μm. **g**. Quantification of shRNA-Control and shRNA-*Pten* organoid morphology upon treatment with DMSO, EZH2 (GSK126) and BAZ2A-BRD inhibitors (BAZ2-ICR and GSK2801). **h**. Representative images of prostate organoid sections immunostained with Ki67.

Next, we analyzed the role of BAZ2A-BRD in PCa initiation using mouse prostate organoids to model PCa driven by loss of *Pten*, one of the most commonly lost tumor suppressor gene in PCa (Berger et al., 2011; Cancer Genome Atlas Research, 2015; Karthaus et al., 2014; Phin et al., 2013; Yoshimoto et al., 2006) (**Fig. 4f-h**). PTEN was also shown to have a pivotal role in CSCs by regulating pathways critical for stem cell maintenance, homeostasis, self-renewal and migration (Luongo et al., 2019; Mulholland et al., 2009; Wang et al., 2006). Furthermore, recent results showed that BAZ2A is required for the initiation of PCa driven by *PTEN*-loss (Pietrzak et al., 2020). Downregulation of *Pten* was achieved by infection of prostate organoids with lentivirus encoding shRNA-*Pten*, which allowed 80% reduction of *Pten* levels (**Supplementary Fig. 4e**). Consistent with previous reports (Karthaus et al., 2014; Pietrzak et al., 2020), compared to control organoids, shRNA-*Pten* organoids displayed altered morphology as evident by the solid and compact structure and elevated expression of nuclear proliferation marker Ki67 (**Fig. 4f-h**). Remarkably, treatment with BAZ2A-BRD inhibitors BAZ2-ICR or GSK2801 impaired *Pten*-loss mediated transformed phenotype as evident by the translucent phenotype, the intact bilayer structure, and low Ki67 signal displayed by the majority of organoids treated with BAZ2A-BRD inhibitors whereas EZH2 inhibitor GSK126 did not show any significant effect. These results are consistent with previous data showing that BAZ2A is required for the initiation of PCa driven by *Pten*-loss (Pietrzak et al., 2020) and highlighted an important role of BAZ2A-BRD in this process. Collectively, these data indicate that pharmacological targeting of BAZ2A-BRD abolish the transition of PCa cells into a CSC-like state and the oncogenic transformation mediated by *Pten*-loss.

## Discussion

Previous work showed that BAZ2A, a component of the chromatin remodeling complex NoRC, is implicated in aggressive PCa (Gu et al., 2015; Pietrzak et al., 2020). In this study, we showed that BAZ2A-BRD is required for the transition of PCa cells into a cancer stem-like state and the initiation of PCa driven by *PTEN*-loss. We showed that BAZ2A-BRD is an epigenetic reader of H3K14ac and is required for the association of BAZ2A with a class of inactive enhancers that are marked by H3K14ac. We showed that the expression of a large fraction of genes in the nearest linear genomic proximity to these BAZ2A-bound enhancers depends on BAZ2A. These genes are implicated in developmental and differentiation pathways, which are usually altered in aggressive and dedifferentiated PCa. The expression analysis of a set of genes that are repressed by BAZ2A and frequently silenced in advanced and poorly differentiated tumors relative to indolent tumors showed the requirement of EP300, one of the writers of H3K14ac. The link of H3K14ac with gene silencing is also consistent with previous studies showing that the transcription regulator ZMYND8, the tumor suppressor PBRM1, and H3K9 methyltransferase SETDB1 can recognize H3K14ac and repress gene transcription (Jurkowska et al., 2017; Li et al., 2016; Liao et al., 2019). The interaction of BAZ2A-BRD with H3K14ac is also of particular interest since H3K14ac was found to be the most prominent histone modification in regulating SNF2H/ISWI nucleosome remodeling and activating only the NoRC complex (Dann et al., 2017). Thus, it is possible that H3K14ac is not only acting for the recruitment of BAZ2A to chromatin but also for the remodeling of nucleosomes that might play a role in the regulation of gene expression. A BAZ2A-mediated remodeling at nucleosomes marked with H3K14ac might therefore be part of the mechanisms to silence gene expression and future studies will address this point.

Cancer stem cells represent a subpopulation within tumors that has an enhanced capacity for therapeutic resistance and thought be implicated in relapse of many tumors types, including PCa (Flemming, 2015; Mayer et al., 2015; Zhang et al., 2017). Thus, efficient targeting of CSCs in cancer treatment is critical for developing effective therapeutics. Cancer cells can transition between stem and differentiated states in response to therapeutic insults or other stimuli within the microenvironment (Plaks et al., 2015; Rich, 2016). In this work, we used an established method to measure the ability of PCa cells to form tumorspheres (Portillo-Lara and Alvarez, 2015; Rajasekhar et al., 2011b). We showed that BAZ2A-BRD is required for the transition of PCa cells into CSCs, suggesting that targeting BAZ2A-BRD with BAZ2A-BRD inhibitors in tumors should be beneficial in impairing the transition of cancer cells toward a stem like-state, thereby decreasing tumor relapse potential.

Current therapeutic approaches for PCa remain insufficient for some patients with progressive disease. Targeting AR axis can manage PCa relapses only for a limited time period since PCas become unresponsive and progress to a castrate-resistant form with limited treatment options (Linder et al., 2018). The development of PCa stem cell-specific anticancer drugs constitutes an attempt to innovate the treatment of PCa (Leao et al., 2017). Previous results showed that AR-signaling-competent human CRPC cell lines are preferentially sensitive to BET inhibitors JQ1 (Asangani et al., 2014). However, inhibition of BET-BRD does not have an effect in AR independent cell lines such as PC3 cells (Asangani et al., 2014). Our study demonstrated that BAZ2A-BRD function can be pharmacologically targeted through BAZ2A-BRD inhibitors in CRPC cells and proposed that inhibition of BAZ2A-BRD can be a potential strategy to use against cancer stem cells in PCa.

## Materials and Method

### Culture of PC3 and PCa spheres

All the indicated cell lines were purchased from the American Type Culture Collection. PC3 cells were cultured in RPMI 1640 medium and Ham’s F12 medium (1:1; Gibco) containing 10% FBS (Gibco) and 1% penicillin-streptomycin (Gibco). DU-145 were cultured in RPMI 1640 medium containing 10% FBS and 1% penicillin-streptomycin. PCa spheres were cultured in serum free DMEM/F12 medium (1:1, Gibco) supplemented with 20ng/ml of basic human FGF (Sigma), 20ng/ml of EGF (Sigma), 3μg/ml of Insulin (Sigma) and 1x B-27 (Gibco). 1×10^4^ cells were seeded in low adhesion petri dishes (Greiner) and cultured for 6 to 8 days. About 20% of media was changed every 2 days. All cells were regularly tested for mycoplasma contamination. PC3 cells were treated with 1μM of BAZ2A-BRD inhibitors BAZ2-ICR (SML1276, Sigma) and GSK2801 (SML0768, Sigma), EZH2 inhibitor GSK126 (S7061-5MG, Lubio), BET inhibitors JQ-1 (SML1524, Sigma) or DMSO as control. Fresh medium supplemented with the drugs was added every 2 days. PCa spheres were culture for 6 days. Cells were collected and transferred to a 24 well plate to image and quantify the number of PCa spheres upon drug treatments.

### Culture of prostate organoids

Isolation of prostate epithelial cells was performed as previously described (Karthaus et al., 2014; Pietrzak et al., 2020). Murine prostate lobes were isolated from 10-11 weeks old mice under dissection microscope, macerated with blade and transferred into tube containing 1.5 ml solution of Collagenase/Hyaluronidase (Stem Cell Technologies) in ADMEM/F12 (Gibco Life Technologies) containing 100 μg/ml Primocin (InvivoGen). Subsequently, prostate tissue was incubated at 37°C rocking for 2 h. After collagenase digestion, samples were centrifuged at 350 g for 5 min, supernatant was discarded and cell pellet was washed with HBSS (Life Technologies). To dissociate prostate tissue into single cells, cell pellet was mixed with 5 ml TrypLE (Gibco Life Technologies) with the addition of 10 μM ROCK inhibitor Y-27632 (STEMCELL Technologies) and incubated for 15 min at 37°C followed by pipetting up and down several times. Subsequently, HBSS + 2% FBS (Gibco) (equal to 2x volume of TrypLE) was added to dissolve TrypLe and quench the reaction. Trypsinized prostate tissue was centrifuged at 350 g for 5 min, supernatant was discarded, and the pellet was washed with HBSS and centrifuged again. 1 ml of pre-warmed Dispase (STEMCELL Technologies)/DNaseI (STEMCELL Technologies) solution was added and samples were vigorously and continuously pipetted up and down until solution was homogenously translucent with no visible tissue fragments. Samples were centrifuged to remove Dispase solution and cell pellet was washed with ADMEM/F12 containing 2% FBS (equal to 5x volume of Dispase). To obtain single cells, suspension was filtered through a 40 um cell strainer, centrifuged and then supernatant was removed. Cells were resuspended in ADMEM/F12 + 2% FBS and viable cells were counted using hemocytometer and trypan blue.Generation of organoids was performed as previously described (Pietrzak et al., 2020). Prostate cells were re-suspended in cold ADMEM/F12 5+ medium at density 5.000 cells per 40 μl for total organoid culture. Cell suspension was mixed with 60 μl of pre-thawed growth factor reduced Matrigel (Corning) and plated in a form of drop into the middle of a well from 24-well plate (TPP). Organoids were cultured in 500 μl of ADMEM/F12 supplemented with B27 (Gibco Life Technologies), Glutamax (Gibco Life Technologies), 10 mM HEPES (Gibco Life Technologies), 2% FBS, 100μg/ml Primocin (ADMEM/F12 5+ medium) and contained following growth factors and components: mEGF (Gibco Life Technologies), TGF-β/Alk inhibitor A83-01 (Tocris), Dihydrotestosterone (DHT) (Sigma-Aldrich) with the addition of 10 μM ROCK inhibitor Y-27632 for the first week after seeding (organoid culture complete medium). Medium was changed every 2-4 days by aspiration of the old one and addition of fresh 500 μl of culture organoid medium. Sequences encoding control shRNA (CACAAGCTGGAGTACAACTAC) and shRNA targeting *Pten* (CGACTTAGACTTGACCTATAT) were cloned into lentiviral vectors. Lentiviral supernatants were concentrated 100 times using Lenti-X Concentrator (Clontech) according to manufacturer’s protocol. 40 μl of concentrated virus was mixed with 100.000 cells suspended in transduction medium (ADMEM/F12, B27 Glutamax, 10 mM HEPES, 100μg/ml Primocin, 2μg/ml Polybrene (Sigma-Aldrich), 10 μM ROCK inhibitor). Cells were incubated with virus for 20 min at RT, centrifuged for 40 min at 800 g at RT, and placed in incubator for 3.5 h to recover. Cells were plated in Matrigel at 5000 cells/well density and after 3 days postseeding 1 μg/ml puromycin (Gibco) was applied to select organoids stably expressing shRNA-control or shRNA-*Pten*.

Organoids were cultured for 22 days in the presence of 5μM of BAZ2A inhibitors BAZ2-ICR (SML1276, Sigma), GSK2801 (SML0768, Sigma), EZH2 inhibitor GSK126 (S7061-5MG, Lubio), BET inhibitors JQ-1 (SML1524, Sigma) or DMSO as control. Medium was replaced every two days with 500μl of fresh medium supplemented with inhibitors.

### Establishment of FLAG-HA-BAZ2A PC3 cell line

In order to introduce the FLAG-HA tag in *BAZ2A* locus a donor plasmid was synthetized by IDT containing the genomic sequence of the FLAG-HA tag flanked by ±1KB homology arm to *BAZ2A* locus. Generation of stable cell was achieved by Alt-R™ CRISPR-Cas9 system from IDT as per manufacturer protocol. A crRNA guide sequence (CCTTCTCTCCCAGTTCTCGG) was chosen to target the *BAZ2A* locus on exon 2 three base pairs upstream of the ATG start codon. Equimolar concentrations of RNA oligos synthetized by IDT (sgRNA and transactivating crRNA) were mixed in Nuclease-Free Duplex Buffer (IDT) to a final concentration of 1μM and were hybridized by heating the mix at 95°C 5min and cooling it down to RT. 1μl of hybridized RNA oligos were incubated with 1μM of Cas9 protein (IDT) in Opti-MEM I Reduced-Serum Medium (Gibco) and incubated at RT for 15min to form a ribonucleoprotein (RNP) complex. RNP complex was transfected for 48h with Lipofectamine RNAiMax (Invitrogen) to 4×10^4^ PC3 previously transfected with donor plasmid carrying the desired inclusion sequence. Single cells clones were isolated and grew until colonies were big enough to be genotyped. To assess the integration of FLAG-HA tag in *BAZ2A* locus, 2 oligos were designed to amplify a genomic region that spans the FLAG-HA tag and a region 150bp outside of the left homology arm that is not encoded in the donor plasmid. Genotyping of clones was performed using primers amplifying the region that spans the insertion site of FLAG-HA tag within *BAZ2A* locus, where a product of 230bp represents the wild type allele and a product of 300bp indicates the integration of the FLAG-HA tag.

### Expression and purification of recombinant proteins

Sequence corresponding to *BAZ2A* bromodomain (residues 1797-1899) was codon optimized for robust expression in E. coli and cloned into pGex4T-1. *BAZ2A* bromodomain mutants (Y1830F, N1873L) were achieved by single point mutation PCR and sequenced to ensure accuracy of the point mutations. Expression of recombinant proteins was achieved by transforming E. coli competent BL21 bacteria with 600ng of plasmids expressing BAZ2A bromodomain. Bacteria was grown at 37°C in Terrific Broth (24g/L yeast extract, 20g/L tryptone, 4mL/L glycerol, 0.017M KH_2_PO_4_, 0.072M K_2_HPO_4_) until reached an OD of 0.6-0.7 at 600nm of absorbance. Induction of expression of recombinant proteins was done at 16°C overnight by supplementing the inoculated culture with 0.5mM IPTG. Bacteria was harvested by centrifugation and pellet was resuspended in 4x (w/v) lysis buffer (20mM Tris pH7.5, 300mM NaCl, 1mM DTT, 20% glycerol, 1mM PMSF) and sonicated 3x 30s on/off. After sonication, lysates were treated with DNase I (50U) and RNase-A (50μg) 30min at 4°C with constant rotation. Lysates were then spun down and supernatant was collected and filtered (0.22μM). GST-fusion proteins were purified using Glutathione-Sepharose 4B beads (GE Healthcare). Purified proteins were eluted with 20mM Gluthathion and dialyzed overnight (20mM Tris pH 7.5, 100mM NaCl, 2mM EDTA, 20% Glycerol). Purity of eluted proteins was assessed by SDS-page.

### Histone peptide arrays

MODified Histone Peptide Arrays were acquired from Active Motif. Briefly, Arrays were blocked for 4h with Odyssey^®^ Blocking Buffer diluted in PBS (1:1). Binding of recombinant proteins to histone peptide array was done by diluting 10nM of recombinant GST or GST-BAZ2A proteins in 3ml of HEMG buffer (25 mM HEPES pH 7.6, 12.5 mM MgCl2, 0.5 m EDTA, 1mM DTT, 0.2 mM PMSF, 10% glycerol) and incubated overnight at 4°C. Pharmacological inhibition of GST-BAZ2A-BRD was achieved by incubating recombinant proteins with 50nM of BAZ2-ICR or GSK2801 overnight in HEMG buffer. Detection of binding was determined by 2h incubation (room temperature) with anti-GST (SC-459, 1:1000 in Odyssey^®^ Blocking Buffer), followed by 1h incubation with goat anti-Rabbit (IRDye^®^ 800CW, 1:10000 in Odyssey^®^ Blocking Buffer). Between incubation periods, histone peptide arrays were washed 3X in PBST to reduce background levels. Arrays were scanned on Odyssey Infrared Imaging System. Active Motif’s Array Analyzer software was used to identify peptides bound by recombinant proteins and intensity of binding was measured as fold changes over background level.

### Plasmid and siRNA transfections

cDNA corresponding to mouse *BAZ2A* wildtype was cloned into AASV1 donor vector (DC-DON-SH01, GeneCopoeia). Site-directed mutagenesis was performed to create *BAZ2A* bromodomain mutant (Y1775F) and sequencing of plasmid was performed to ensure fidelity of sequences. 1×10^6^ of PC3 cells were transfected for 48h with 8ug of plasmids with X-tremeGENE HP DNA Transfection Reagent (Roche). siRNA against *BAZ2A* (Cat. # SI00102389) and siRNA control (Cat. # 1027281) were obtained from Qiagen. EP300 siRNA (Cat. # J-003486-11-0002) and BAZ2B siRNA (Cat. # D-020487-02-0005) was obtained from Dharmacon. Transfections of siRNA were performed with Lipofectamin RNAiMax (Invitrogen) in Opti-MEM Reduced-Serum Medium (Gibco).

### Chromatin immunoprecipitation

ChIP experiments were performed as previously described (Leone et al., 2017). Briefly, 1% formaldehyde was added to cultured cells to cross-link proteins to DNA. Isolated nuclei were then lysed (50 mM Tris-HCl pH 8.1, 10 mM EDTA and 1% SDS) and sonicated using a Bioruptor ultrasonic cell disruptor (Diagenode) to shear genomic DNA to an average fragment size of 200 bp. 20 μg of chromatin was diluted tenfold with ChIP buffer (16.7 mM Tris-HCl pH 8.1, 167 mM NaCl, 1.2 mM EDTA, 0.01% SDS and 1.1% Triton X-100) and precleared for 2h with 10ul of packed Sepharose beads for at least 2h at 4°C. Immunoprecipitation was done overnight with the indicated antibodies. Pulldowns were done with Dynabeads protein-A (or - G, Millipore) for 4h at 4°C. For HA-BAZ2A ChIPs, preclearing and pulldowns were done with beads block with BSA (50μg) and ytRNA (50μg), per 400ul of Dynabeads protein-A. After washing, elution and reversion of cross-links, eluates were treated with RNAse A (1μg). DNA was purified with phenol-chloroform, ethanol precipitated and quantified by quantitative PCR.

### ChIP-seq analysis

The quantity and quality of the isolated DNA was determined with a Qubit® (1.0) Fluorometer (Life Technologies, California, USA) and a Bioanalyzer 2100 (Agilent, Waldbronn, Germany). The NEBNext^®^ Ultra™ II DNA Library Prep Kit from NEB (New England Biolabs, Ipswich, MA) was used in the succeeding steps. ChIP samples (1ng) were end-repaired and adenylated. Adapters containing the index for multiplexing were ligated to the fragmented DNA samples. Fragments containing adapters on both ends were enriched by PCR. The quality and quantity of the enriched libraries were measured using Qubit® (1.0) Fluorometer and the Tapestation (Agilent, Waldbronn, Germany). The product is a smear with an average fragment size of approximately 300 bp. The libraries were normalized to 10nM in Tris-Cl 10 mM, pH8.5 with 0.1% Tween 20. The TruSeq SR Cluster Kit v4-cBot-HS (Illumina, Inc, California, USA) was used for cluster generation using 8 pM of pooled normalized libraries on the cBOT. Sequencing was performed on the Illumina HiSeq 2500 single end 125 bp using the TruSeq SBS Kit v4-HS (Illumina, Inc, California, USA). Obtained ChIPseq reads from Ilumina HiSeq 2500 were aligned to the human hg38 reference genome using Bowtie2 (version 2.2.5). Read counts were computed and normalized using “bamCoverage” from deepTools (version 2.0.1; (Ramirez et al., 2014)) using a bin size of 50bp. deepTools was used to generate all heat maps, Pearson correlation plots. Peaks from ChIPseq experiments were defined using MACS2 (version 2.1.0; (Zhang et al., 2008)) comparing ChIP signal against input. For all histone modifications default settings were used. BAZ2A peaks were determined using a p-value cutoff as <0.01. Data sets from H3K4me1 (GSM2534170), H3K9me2 (GSM2534657), H3K9me3 (GSM2533935), DNAseq (GSM2400265) were obtained from ENCODE and processed as previously described. Integrative Genome Viewer (IGV, version 2.3.92 (Robinson et al., 2011) was used to visualize and extract representative ChIPseq tracks. Genomic annotation of ChIPseq peaks was done with ChIPseeker (Yu et al., 2015), and promoter regions were defined as ±5Kb from TSS. Analysis of overlapping genomic regions was done with bedtools (version 2.27.1 (Quinlan and Hall, 2010)). Enhancer coordinates were obtained from GeneHancer V4.4 (Plaschkes et al., 2017) and the read coverage +/- 1Kb from the center of peak was obtained using “computeMatrix” from deeptools with a bin size of 50bp. Mean read coverage was then used to classify the regions based on presence or absence of H3K27ac and H3K4me1.

### RNA extraction, reverse transcription, and quantitative PCR

RNA was purified with TRIzol reagent (Life Technologies). 1µg total RNA was primed with random hexamers and reverse-transcribed into cDNA using MultiScribe™ Reverse Transcriptase (Life Technologies). Amplification of samples without reverse transcriptase assured absence of genomic or plasmid DNA (data not shown). The relative transcription levels were determined by normalization to *GAPDH* mRNA levels. qRT-PCR was performed with KAPA SYBR^®^ FAST (Sigma) on Rotor-Gene RG-3000 A (Corbett Research).

### RNAseq and data analysis

Total RNA from 2 independent samples form PC3 cells and PCa cells, and from 3 independent experiments on PC3 cells treated with siRNA control or siRNA*BAZ2A*, were purified with TRIzol reagent (Life Technologies) as stated above. DNA contaminants were removed by treating RNA with 2U Turbo-DNaseI (Invitrogen) for 1h at 37°C and the RNA samples were re-purified using TRIzol. The quality of the isolated RNA was determined by Bioanalyzer 2100 (Agilent, Waldbronn, Germany). Only those samples with a 260nm/280nm ratio between 1.8–2.1 and a 28S/18S ratio within 1.5–2 were further processed. The TruSeq RNA Sample Prep Kit v2 (Illumina, Inc, California, USA) was used in the succeeding steps. Briefly, total RNA samples (100-1000ng) were poly A enriched and then reverse-transcribed into double-stranded cDNA. The cDNA samples was fragmented, end-repaired and polyadenylated before ligation of TruSeq adapters containing the index for multiplexing Fragments containing TruSeq adapters on both ends were selectively enriched with PCR. The quality and quantity of the enriched libraries were validated using Qubit® (1.0) Fluorometer and the Caliper GX LabChip® GX (Caliper Life Sciences, Inc., USA). The product is a smear with an average fragment size of approximately 260bp. The libraries were normalized to 10nM in Tris-Cl 10mM, pH8.5 with 0.1% Tween 20. The TruSeq SR Cluster Kit HS4000 (Illumina, Inc, California, USA) was used for cluster generation using 10pM of pooled normalized libraries on the cBOT. Sequencing was performed on the Illumina HiSeq 2500 single end 100bp using the TruSeq SBS Kit HS2500 (Illumina, Inc, California, USA). Reads were aligned to the reference genome (hg38) with Subread (i.e. subjunc, version 1.4.6-p4; (Liao et al., 2013)) allowing up to 16 alignments per read (options: –trim5 10 –trim3 15 -n 20 - m 5 -B 16 -H –allJunctions). Count tables were generated with Rcount (Schmid and Grossniklaus, 2015) with an allocation distance of 10bp for calculating the weights of the reads with multiple alignments, considering the strand information, and a minimal number of 5 hits. Variation in gene expression was analyzed with a general linear model in R with the package edgeR (version 3.12.0; (Robinson and Oshlack, 2010)) according to a crossed factorial design with two explanatory factors (i) siRNA against *BAZ2A* and a mock sequence and (ii) expression pattern of PCa spheres versus PC3 cells. Genes differentially expressed between specific conditions were identified with linear contrasts using trended dispersion estimates and Benjamini-Hochberg multiple testing corrections. Genes with a *P-*value below 0.05 and a minimal fold change of 1.5 or 2 were considered to be differentially expressed. These thresholds have previously been used characterizing chromatin remodeler functions (de Dieuleveult et al., 2016). Gene ontology analysis was performed with David Bioinformatics Resource 6.8 (Huang et al., 2009).

### Gene Set Enrichment Analyisis

GSEA (Subramanian et al., 2005) were obtained by comparing expression values of PCa spheres and PC3 cells of genes upregulated in PCa spheres (*P* < 0.05, log2FC>1). GSEA *P* value was calculated over 100000 permutations based on the phenotype and FDR was <25%.

### Immunohistochemistry

Organoids were fixed using 4% paraformaldehyde (PFA) overnight and then washed with 70% ethanol. Fixed organoids were pre-embedded in HistoGel (Thermo Fisher Scientific) to pellet organoids together. Tissues and organoid pellets embedded in HistoGel were processed with an automated tissue processor and subsequently embedded in paraffin wax according to standard techniques. Samples were sectioned at 4 μm onto SuperFrost^®^ Plus (Thermo Scientific) microscope slides and air-dried 37 °C overnight. Immunohistochemistry was performed using BrightVision+ histostaining kit (Immunologic) according to manufacturer’s protocol. Briefly, organoid sections were dewaxed in Xylene (3x 3 min), followed by rehydration steps in descending percentages of EtOH (3 min in 100%, 95%, 90%, 70%, 50%), and washed twice in PBS. Slides were boiled in sodium citrate pH 6.0 (10mM sodium citrate, 0.05% Tween 20) for 15 minutes and then allowed to cool to room temperature for 30 minutes. Slides were then washed in PBS and blocked for at least 1 hour in PBS + 2% BSA. Samples were incubated with anti-Ki67 primary antibody (1:100, Abcam ab15580) overnight in blocking reagent at 4°C and then washed 3 times with PBS. Post-antibody Blocking solution was added for 15 min and samples were incubated at RT. Sections were washed twice with PBS and then were incubated for 30 min at RT with Poly-HRP-Goat anti Mouse/Rabbit IgG. After incubation sections were washed twice with PBS and were incubated with 3,3’ Diaminobenzidine (DAB) mixture (Bright-DAB, Immunologic) for 8 min according to manufacturer protocol. After incubation organoid sections were rinsed in deionized water and mounted with Fluoro-Gel.

## Accession numbers

All raw data generated in this study using high throughput sequencing are accessible through NCBI’s GEO (GSE131268).

## Acknowledgements

We thank Dominik Bär for technical assistance and Rostyslav Kuzyakiv for help in bioinformatic analyses. We thank Catherine Aquino and the Functional Genomic Center Zurich for the assistance in sequencing. This work was supported by the Swiss National Science Foundation (310003A-152854 and 31003A_173056), the National Center of Competence in Research RNA & Disease (funded by the SNSF), Sassella Stiftung (to SF and KP), Novartis, Julius Müller Stiftung, Olga Mayenfisch Stifung, Krebsliga Zurich, Krebsliga Schweiz (KFS-3497-08-2014 and KLS-4527-08-2018).

## Supplementary Figures

**Supplementary Figure 1.**
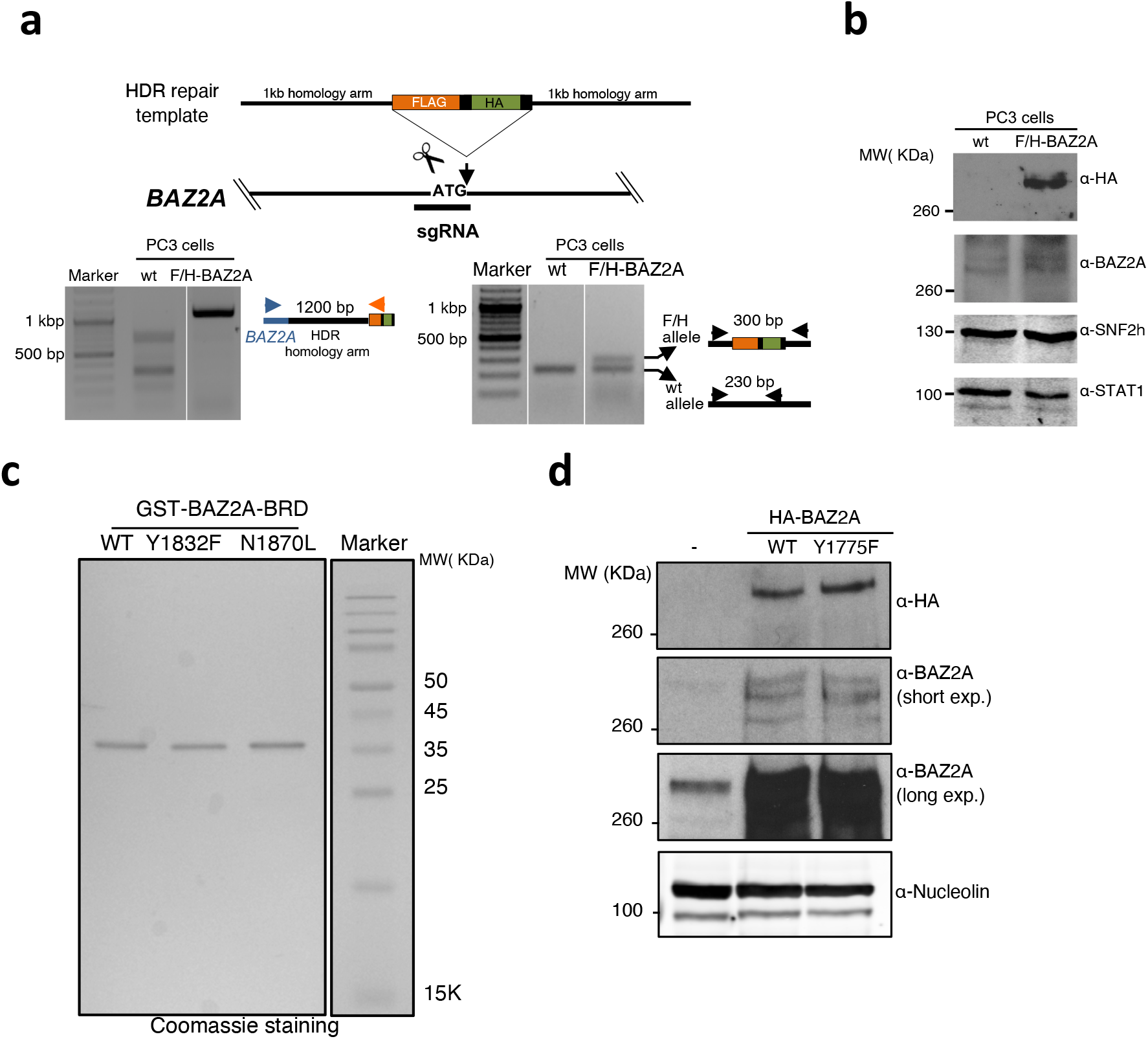
**a**. Schematic representation of the strategy to generate PC3 cell line with FLAG-HA (F/H) tag insertion at *BAZ2A* locus by CRISPR/Cas9 method. Lower panel. PCR genotyping to measure the insertion of F/H sequences into *BAZ2A* locus.. Arrows represent the primers used for PCR genotyping. **b**. Immunoblot of PC3 wild-type (wt) cells and the PC3 cell line expressing endogenous BAZ2A with F/H tag showing comparable levels of BAZ2A. SNF2h and STAT1 serve as protein loading control. **c**. Representative protein gel showing the purity of recombinant GST-BAZ2A-BRD wild type (WT)) and mutants (GST-BAZ2A-BRD_Y1830F_ or GST-BAZ2A-BRD_N1873L_). Proteins are visualized by Coomassie staining. **e**. Western blot showing equal expression of ectopically expressed HA-BAZ2Awt and HA-BAZ2A-BRD mutant (Y1775F) in transfected PC3 cells and endogenous levels of BAZ2A in untransfected PC3 cells. Whole cell lysates from equivalent amounts of cells were analyzed. Nucleolin is shown as a protein loading control.

**Supplementary Figure 2.**
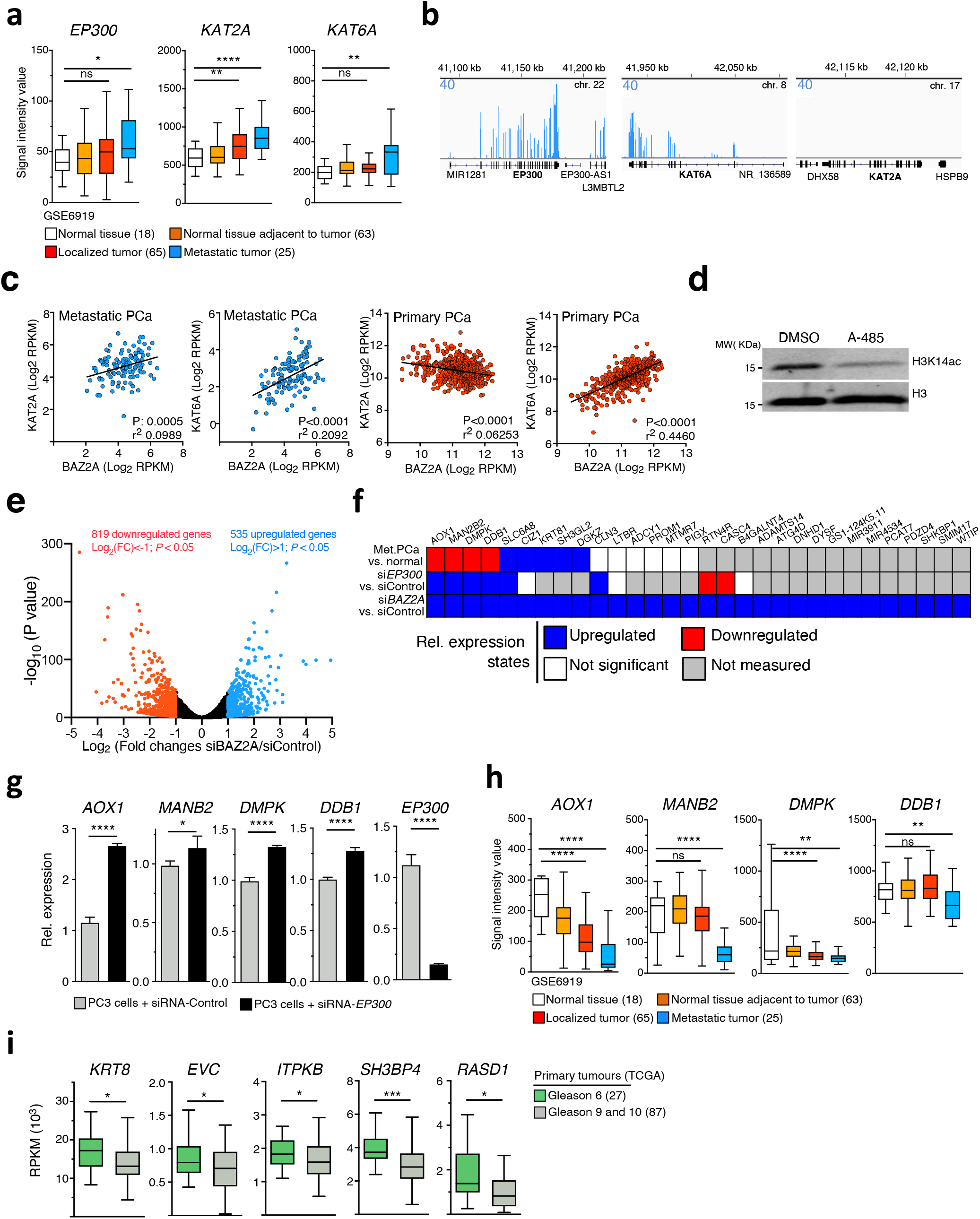
**a**. The H3K14ac histone acetytransferases *EP300, KAT2A*, and *KAT6A* are highly expressed in metastatic prostate cancer tissues. Boxplots showing expression profiles from Gene Expression Omnibus (GEO) data sets GSE6919 (gene expression microarray data) (Chandran et al., 2007; Yu et al., 2004). *< 0.05, **< 0.01, \*\*\*\**P* < 0.0001, two-tailed Student’s *t* test; ns, not significant. **b**. Wiggle tracks showing RNAseq expression profiles of *EP300, KAT2A*, and *KAT6A* in PC3 cells. *KAT2A* is not expressed in PC3 cells due to deletion of the entire locus. **c**. Scatter plot showing the expression of *KAT2A* and *KAT6A* vs. BAZ2A in two large cohorts of metastatic and primary PCa. Data were from (Cancer Genome Atlas Research, 2015; Robinson et al., 2015). **d**. EP300 acetylates H3K14 in PC3 cells. Western blot showing H3K14ac levels in PC3 cells treated for three days without and with the selective catalytic EP300 inhibitor A-485 (10 μM). Histone H3 serves as protein loading control. **e**. Volcano plot showing fold change (log_2_ values) in transcript levels of PC3 cells upon BAZ2A knockdown. Gene expression values of two replicates were averaged and selected for 1.5 fold changes and *P* <0.05. **f**. Table showing expression changes of genes with BAZ2A-bound promoter upregulated upon BAZ2A-KD in PC3 cells in metastatic PCa vs. normal prostate tissue and in PC3 cells depleted of EP300 via siRNA. **g**. qRT-PCR showing increased expression levels of genes with BAZ2A-bound promoter and downregulated in metastatic tumors (*AOX1, MANB2, DMPK*, and *DDB1* - see **F**) in PC3 cells treated with siRNA-EP300. Data were from three independent experiments. Values were normalized to *GAPDH* mRNA. Statistical significance (P-values) was calculated using two-tailed t-test (*<0.05, ****<0.0001). **h**. Boxplots showing expression profiles of *AOX1, MANB2, DMPK*, and *DDB1* from GEO data sets GSE6919 (gene expression microarray data) (Chandran et al., 2007; Yu et al., 2004). **< 0.01, \*\*\*\**P* < 0.0001, two-tailed Student’s *t* test; ns, not significant. **i**. Boxplots showing expression of BAZ2A-bound C2-enhancer genes KRT8, *EVC, ITPKB, SH3BP4*, and *RASD1* in dedifferentiated and aggressive primary tumors scored with Gleason 9 and 10 and indolent tumors assigned with Gleason 6. Data are from TCGA_PRAD (Cancer Genome Atlas Research, 2015).

**Supplementary Figure 3.**
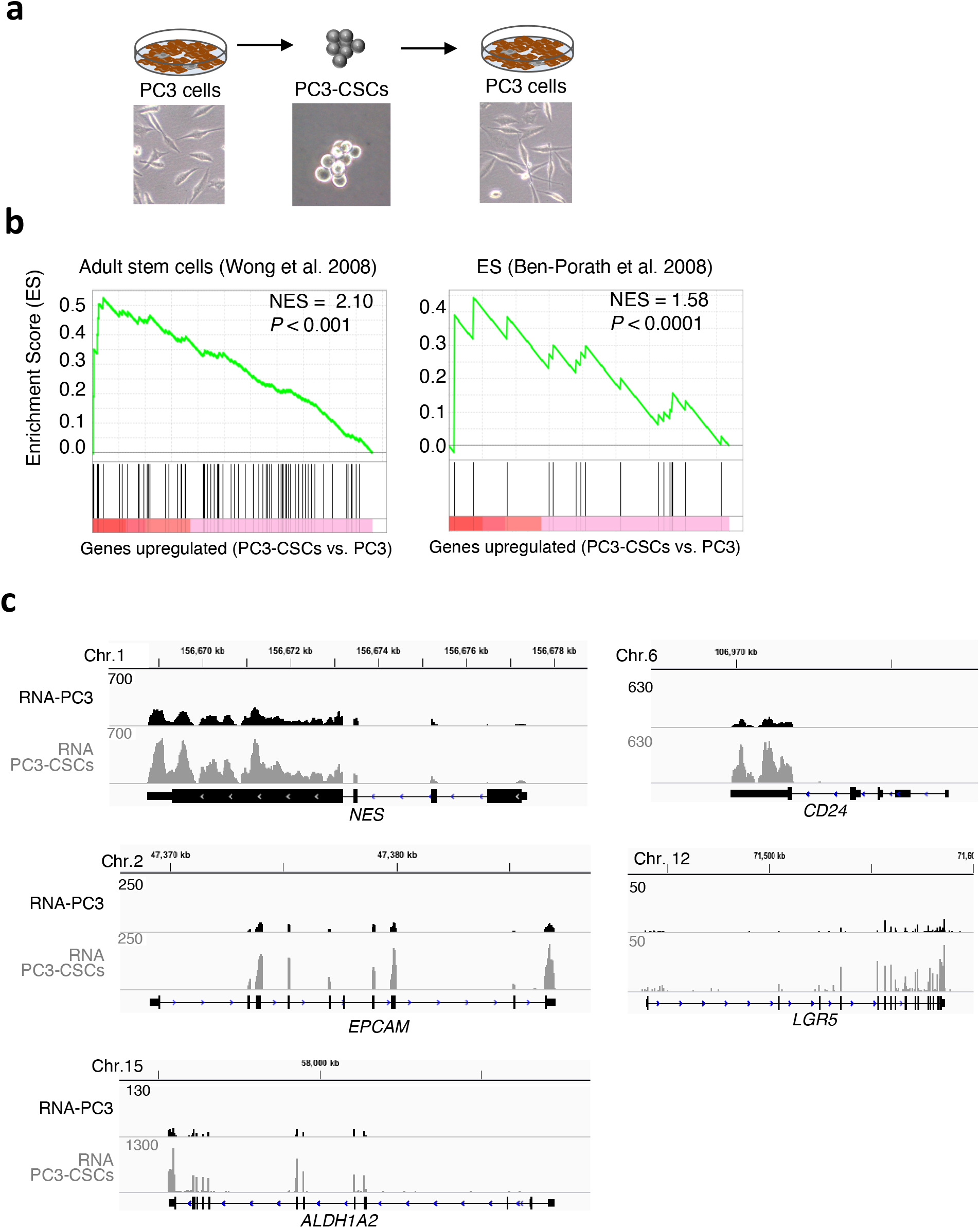
**a**. Representative images showing PC3 cells, the derived tumorspheres PC3-CSCs and PC3 cells generated from PC3-CSCs under adherent cell culture conditions. **b**. Genes upregulated in PC3-CSCs are enriched in stem cell-like signature. GSEA analysis was obtained by comparing expression values of tumorspheres vs. PC3 cells of genes upregulated in PC3-CSCs (*P* < 0.05, log_2_FC>1). *P* was calculated over 100000 permutations based on the phenotype and FDR was < 25%. NES: normalized enrichment score. **c**. Representative wiggle tracks from RNAseq of PC3 and PC3-CSCs of cancer stem cell markers that are upregulated in PC3-CSCs compared to the parental PC3 cells.

**Supplementary Figure 4.**
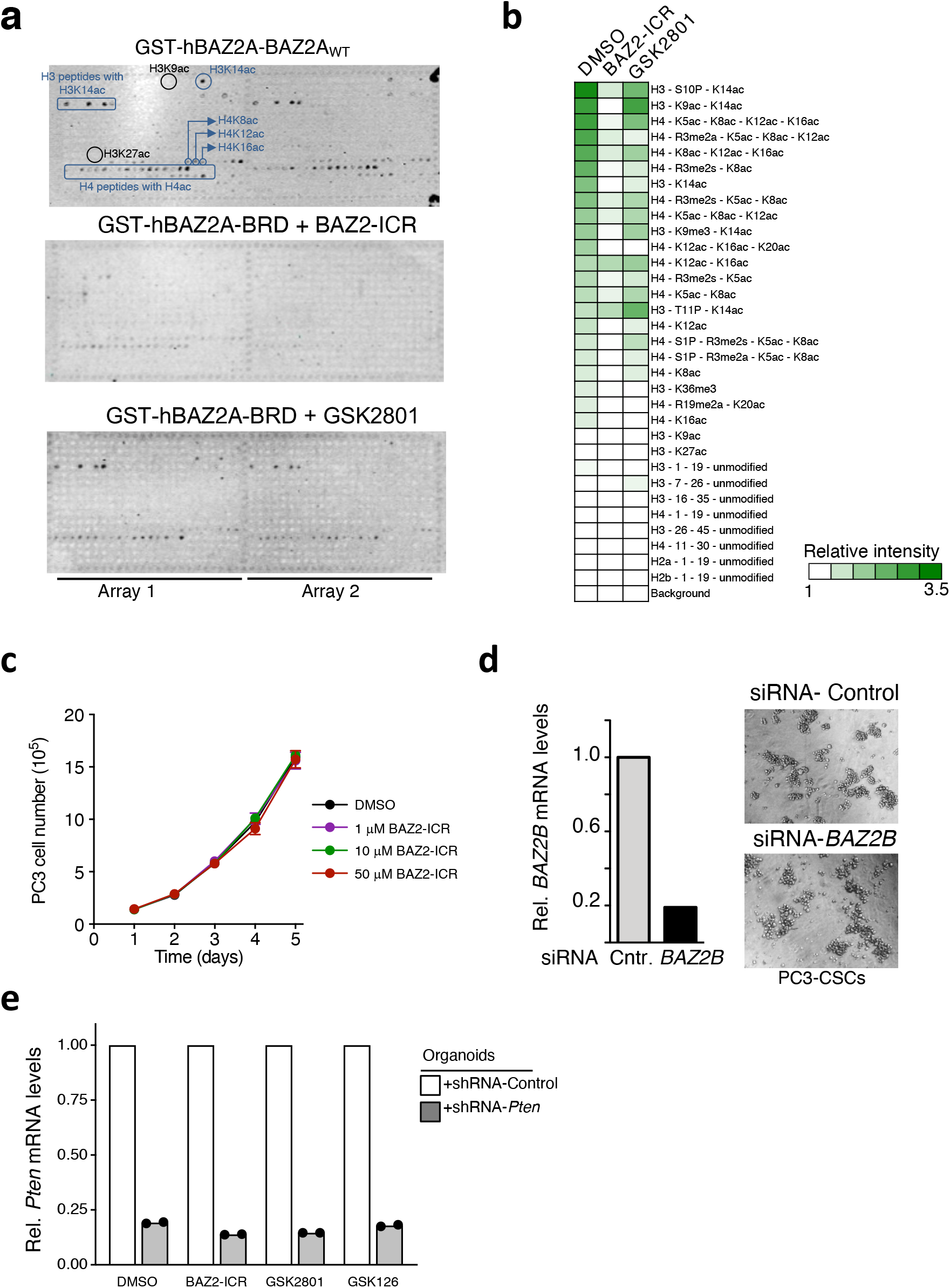
**a**. Images of histone peptide arrays incubated with 10nM of recombinant GST-BAZ2A-BRD_wt_ in the presence of 50nM of BAZ2A-BRDi, BAZ2-ICR or GSK2801. Visualization of binding was performed by incubation with anti-GST antibodies and imaged on Odyssey Infrared Imaging System. For a better visualization of the data, the image of GST-BAZ2A-BRD_wt_ without BAZ2A-BRDi shown in **Figure 1D** has been included. Blue circles mark peptides recognized by BAZ2A-BRD (i.e. H3K14ac), whereas black circles show some of the acetylated peptides not recognized by BAZ2A-BRD (H3K27ac and H3K9ac). **b**. Heatmaps showing the relative BAZ2A-BRD binding intensity at modified histones peptides upon incubation with BAZ2A-BRDi BAZ2-ICR or GSK2801. Binding intensity was calculated as average of fold change from peptides signal over background controls of 2 different arrays. **c**. Cell proliferation curves of PC3 cells treated with 5, 10, and 50μM BAZ2-ICR. Error bars represent the standard deviation from three independent experiments. **d**. *BAZ2B* depletion in PC3 cells does not affect the dedifferentiation into PC3-CSCs. Left panel: *BAZ2B* mRNA levels were measured by qRT-PCR in PC3-CSCs and normalized to *GAPDH* mRNA. Right panel: Representative figures showing PC3-CSCs generated from PC3 cells transfected with siRNA-Control or siRNA-*BAZ2B*. **e**. qRT-PCR showing *Pten* mRNA levels in mouse prostate organoids expressing shRNA-control and shRNA-*Pten* and treated with DMSO, BAZ2-ICR (5μM), GSK2801 (5μM) or GSK126 (5μM). Values are from experiments. Data were normalized to *Hprt* mRNA and organoids treated with siRNA-Control.

## Notes

### Competing Interest Statement

The authors have declared no competing interest.

